# BarlowTrack: A Self-Supervised Framework for Zero-Shot Multi-Object Cell Tracking

**DOI:** 10.1101/2025.11.10.687595

**Authors:** Charles Fieseler, Itamar Lev, Jalaja Madhusudhanan, Zihao Zhai, Siegfried Schwartz, Manuel Zimmer

## Abstract

Recent advances in neuroscience have made it possible to image large brain regions at single-cell resolution. However, classic methods for processing these videos into neuronal time series fail in the presence of large and nonrigid deformations, in particular for freely moving animals. Several successful algorithms have been proposed to solve this problem in moving *C. elegans*, but they are highly specific to the conditions of a single research setting. We propose a tracking pipeline based on self-supervised learning that achieves a high level of zero-shot accuracy across conditions, and, for the first time, independent laboratories. We contribute a novel term in the Barlow Twins loss function to encourage decorrelation of features across detected instances at the same time point. To encourage broad adoption, we use the standardized Neurodata Without Borders (NWB) format and we provide a GUI for visualization of the final results and neuronal time series. Finally, we provide a benchmark of datasets with ground truth annotations in the NWB format for further algorithmic development.

## 1 Main

Many questions in modern neuroscience require simultaneous access to natural behavior and neuronal data, especially at single-cell resolution. This has been possible using head-fixed monkeys^1^, drosophila^2,3^, mouse^4^ and other organisms for many years; systems that record more natural behavior but with a limited field of view have also been developed, especially in mouse^5,6^. Systems with both free motion and that can record all or nearly all of the brain have also recently been developed in zebrafish larvae^7^, hydra^8^, and *C. elegans*^9–11^. This freely moving setting is challenging due to the large and nonrigid deformations that are generated during natural motion, and can be broken down into two sub-problems. The first is cell segmentation, for which there are many general solutions^12–15^ and for which foundation models have recently been developed that can succeed across imaging modalities^16,17^. Although segmentation is still an active area of research, accurate performance can be achieved via published methods. The second subproblem, tracking or associating objects across time, cannot be solved with general published algorithms. For this reason, many labs have devoted years to developing organism-specific and lab-specific solutions, which, while impressive, lack generalizability across conditions.

In the *C. elegans* community alone, 6 labs have published 9 unique pipelines to perform whole-brain freely moving *C. elegans* imaging^9,10,18–23^ including completely manual tracking requiring 200 hours to annotate a 10 minute video^24^; several other approaches have been published for the semi-immobilized case^25–27^. The diversity of approaches demonstrates that this is not a solved problem and for each new set of conditions, and in particular for a different scientific lab, the problem must be solved again.

For image processing tasks, supervised learning via Convolutional Neural Networks (CNNs) and more recently Vision Transformers (ViTs) dominate benchmarking challenges, including cell tracking^28^ and segmentation^17^. However, the need for ground truth annotation is a major bottleneck in generalization. For this reason, self-supervised learning was developed^29^, which is a training paradigm that allows a neural network to learn a feature space without ground truth annotations. In this paper, we use a novel modification of a recently developed self-supervised learning architecture (Barlow Twins^30^) in order to learn a feature space that separates each segmented neuron, allowing a simple clustering approach to connect all objects across time. We show that this method generalizes both across conditions and across labs, the first algorithm to do so. We further developed a GUI for data visualization as well as a python package for user-friendly analysis of new data. This GUI allows reproducible assignment of neuronal IDs based on anatomy and time series activity patterns. These packages are compatible with the Neurodata Without Borders format, which has recently been adopted by the *C. elegans* community^31^, and we release several previously published datasets in this format. Taken together, our solution both solves a longstanding problem and provides benchmark datasets for new solutions to be developed.

## 2 Results

### 2.1 Motivation and Problem Context

Tracking neurons in freely behaving animals is a challenging but highly structured task. Natural locomotion is nearly sinusoidal (1a), and ≈ 95% of the variance can be modeled with only 4 PCA components^32^. Due to this low dimensionality, one might expect that the neuron tracking problem could be solved with off-the-shelf point cloud matching algorithms. However, generic methods have been shown to have very poor performance, with custom methods performing much better^22^. Neurons are densely packed within the head region (1b), and even small tracking inaccuracies could cause cascading identification problems. Thus, we first sought to quantify the motion of single neurons at a higher resolution.

The motion across all neurons can be visualized as vector fields or color-coded flow diagrams (1c), showing some simple rotations (first panel), as well as nonrigid compression of the nose (second panel). Indeed, the displacement between adjacent time points (0.3 seconds for the Zimmer lab datasets; see Table 1) is often much larger than the distance to the nearest neighbor (1c). This is true even though the motion in the image is less than the physical motion of the animal; the camera follows the worm, centering it and reducing centroid motion to near 0. Quantifying the complexity of the motion is not trivial, but like the overall body posture, the displacement vector field of individual objects can be modeled using Principle Component Analysis (PCA); specifically, we use probabilistic PCA due to the missing values^33^. The neuron vector field is much higher dimensional, with ≈ 20 modes needed to capture 90% of the variance (1d red line). However, pure variance explained is not the same as the tracking task, and in particular denser regions require smaller errors in order to achieve proper identification. Thus we also quantified the single-step tracking accuracy, i.e. how many neurons will be closer to their ground truth position than to their nearest neighbor (See methods for more detail). This metric, gives a similar accuracy as the variance explained, with 20 modes giving 90% tracking accuracy at the next time point. For many tasks this accuracy may not be sufficient, and to achieve a higher accuracy of 95% many more modes (around 50) are needed. Thus the motion at the single-cell level creates a structured but high dimensional tracking problem.

**Table 1.** Overview of datasets and physical resolutions. Dataset names use the name of the PI (Manuel Zimmer, Steven Flavell, and Andrew Leifer). # objects means the number of objects in the ground truth for the datasets that have ground truth. *means that the ground truth of the Flavell lab datasets is not manually annotated, but rather the tracking is from an algorithm designed specifically for those datasets.

| Dataset name | Resolution<br>$\mu\text{m}/\text{pixel}$ | VPS<br>(Hz) | #animals | #volumes | #objects | Manual GT? |
| --- | --- | --- | --- | --- | --- | --- |
| Zimmer<sup>10</sup> | 0.35x0.35x1.5 | 3.5 | 3 | 1667 | 60-130 | Yes |
| Zimmer 2 (This work) | 0.35x0.35x1.5 | 3.5 | 5 | 1000-1667 | 100-180 | No |
| Flavell<sup>9</sup> | 0.33x0.33x0.33 | 1.7 | 5 | 3200 | 130-180 | No* |
| Leifer<sup>18</sup> | 0.32x0.32x1.5 | 6 | 1 | 1536 | 79 | Yes |

Significant progress has been made in automating this problem^9,10,18–23^, but no methods have been successfully applied across scientific labs. This is somewhat surprising, because all imaging pipelines are similar in the *C. elegans* community. On the genetic side, current methods augment the calcium sensitive GCaMP channel with asecond reference channel that is stable across time; both markers are generally localized to the cell nucleus. On the microscopy side, all current methods use confocal spinning-disk fluorescence microscopy; light sheet microscopy technologies are being developed^34,35^ but there is no published work applying them to freely moving animals. Note that although tracking in all published methods is performed exclusively on the reference channel, segmentation may be performed on both.

Regardless of these basic similarities, many conditions will lead to dramatic quantitative and qualitative differences in generated images. In particular, several factors directly influence the motion experienced between volumes, namely the spatial and temporal resolutions, as well as the speed of the animal itself. The volume rate of data collection varies significantly across labs (from 1.7Hz^9^ to 10Hz^24^), as does the spatial resolution particularly in the z axis (from 0.33 µm^9^ to 1.5 µm^10^; see Table 1). The speed of the animal, and thus inter-volume motion of neurons, is strongly influenced by genotype^36^, the density of the material on which the worms crawl^37^, and presence of food or other sensory stimuli^38,39^. Further changes in the image can be caused by different promoters (genetic driver for the reference channel) or the expression of additional genetic markers. The presence of other animals, necessary for studies of social or mating behaviors^24^, can also dramatically change the local neighborhood of single neurons and the global image features. Progress in solving these differences is further hindered by incompatible and often undocumented data formatting.

### 2.2 Data preparation: Neurodata Without Borders Format and Cell Detection

Standardizing data formats across scientific projects is not a new problem. The Neurodata Without Borders (NWB) format^40^ is the outcome of a major organizational effort across the field of neuroscience to specify a data format that allows all elements of an experiment to be contained in one file, from raw imaging data and intermediate object segmentation, to final production of time series. Recently, this format has been extended for the volumetric whole brain freely moving case in *C. elegans*^41^, but no published pipelines for freely moving animals are compatible with it (with the exception of our prior work^10^, which has limited generalization performance; see Table 2). However, many datasets across different labs are being formatted in this way, and it is becoming the standard^31,41^.

**Table 2.** Benchmark results including other tracking algorithms; accuracy for BarlowTrack networks is reported as the median accuracy of networks trained with default settings. * means that the algorithm requires partial ground truth annotations in order to produce tracks.

| Algorithm name | Zimmer | Flavell | Leifer |
| --- | --- | --- | --- |
| NeRVe<sup>18</sup> | — | — | 82.9% |
| fDNC<sup>22</sup> | — | — | 79.1% |
| CeNDeR<sup>19</sup> | — | — | 88.25%* |
| ZephIR<sup>20</sup> | — | — | 94.5%* |
| Zimmer Tracklet Method<sup>10</sup> | 91.4% | 3.5% | — |
| BarlowTrack (HDBSCAN) | 96.7% | 94.4% | 96.2% |
| BarlowTrack (Label Propagation) | **97.3%** | **96.7%** | **97.4%** |

Next, the first image processing problem must be solved: object detection or instance segmentation. There are many published pipelines to solve this problem for single cell-resolution microscopy, and in the *C. elegans* community most utilize one of two approaches. The first is to perform foreground-background separation using a threshold or U-net architecture, then detect peaks, and finally generate objects using a simple radius^24^ or a watershed algorithm^18^. The second strategy is to perform instance segmentation in an end-to-end manner using a trained neural network^9,10,23^, for example using the StarDist^12,13^ architecture. For example, all Zimmer lab datasets used the same pretrained StarDist model using *≈* 10 partially annotated ground truth volumes.

### 2.3 Overview of BarlowTrack: Tracking via self-supervised neural network

Tracking consists of two steps: (1) embedding 3d-cropped neurons in a feature space (Fig. 2a-b), and (2) clustering that feature space (Fig. 2c). The feature space is trained using self-supervised learning, removing the need for any manual annotation. We present a novel time-aware clustering algorithm (Section 2.3.2), but it can also be performed using optimized off-the-shelf code implemented in Python (such as HDBSCAN^42^; see Fig. S1c).

#### 2.3.1 Training a feature space network

We based our network on the encoder half of the Residual 3d-Unet architecture^43^, specifically three Conv3D layers with LeakyReLU activation, followed by MaxPool3d layers. The projection head is a single layer feedforward network with sigmoid activation, followed by a linear layer.

We train this network using self-supervised learning, implementing a modified Barlow Twins loss function^30^. The original loss function balances an invariance term and a Redundancy Reduction (RR) term:

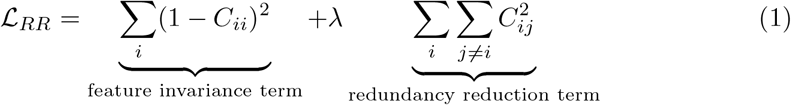

Where the *C*_*ii*_ and *C*_*ij*_ terms are the correlation matrix in the feature space, calculated across the batch:

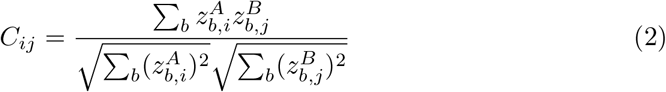

where b indexes batch samples while i and j index the feature space, which is the network’s output. C is a square matrix with size equal to the embedding dimension hyperparameter, and with absolute values between 0 (no correlation) and 1 (perfect correlation or anti-correlation).

We augment this classic loss with the same cross-correlation loss calculated not across the batch, but across the features. This gives rise to the Object Space (OS) loss term:

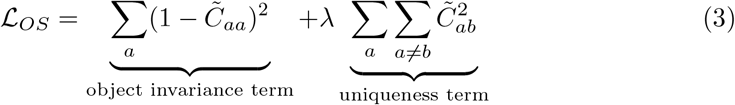

Where the 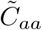 and 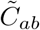 terms are the correlation matrix in the *batch* space, calculated across the *features*:

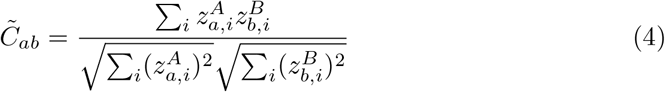

The original invariance term, which encourages individual features *across objects* in the embedding space to be invariant to augmentations. This invariance term does the reflection of that: it encourages the features *of individual objects* in the batch to be invariant to those same augmentations. However, the uniqueness term is quite different, and encourages the embeddings of individual objects to have zero correlation. This assumption is not satisfied in the general image processing case because different objects in a batch may have the same identity or class, although even treating all samples as different can be beneficial^44^. However, in the object tracking case, each object in a time point must be different from all others, easily satisfying the core assumption.

Thus, this loss function requires this restriction on how batches are chosen: objects from the same time point. In this study and for the number of neurons in *C. elegans* (302 neurons in the entire brain), we set the batch equal to all objects found in a time point.

#### 2.3.2 Tracking via clustering

After training a BarlowTrack network to produce embeddings, the objects must be associated across time. This can be done via direct clustering in the embedding space. If the embedding space solves the tracking problem well, then the embedding of the same object at all time points will be more similar than the embeddings of other objects. This can be visualized by the clear separation of objects in a 2d UMAP projection (Fig. 2), which are generally assigned to the same color (cluster). However, even with the same trained network, the exact clustering method will change the quality of the final tracks.

We benchmarked several different clustering approaches, including the unsupervised HDBSCAN algorithm. However, because it is unsupervised, HDBSCAN occasionally completely missed neurons which formed lower density clusters in the embdedding space, marking them as outliers. Thus we designed a custom clustering algorithm and found that it performed better across all ground truth datasets (Table 2; Fig. S1c). Our method uses the knowledge that at a single time point, all objects are distinct. Thus, after constructing a nearest-neighbor graph in embedding space, label propagation^45,46^ is run from seeded points, corresponding to objects at one time point. Because these seeds form a very sparse set of labels, we clamp the information on those nodes instead of the more common averaging. The seed index at different time points is meaningless and therefore multiple seeded runs (labelings) must be aligned (relabeled); for this, we use a custom time-aware implementation of spectral ensemble clustering (similar to^47^). Psuedocode for the clustering pipeline is outlined in Alg. 1.

##### Algorithm 1

Time-aware Clustering

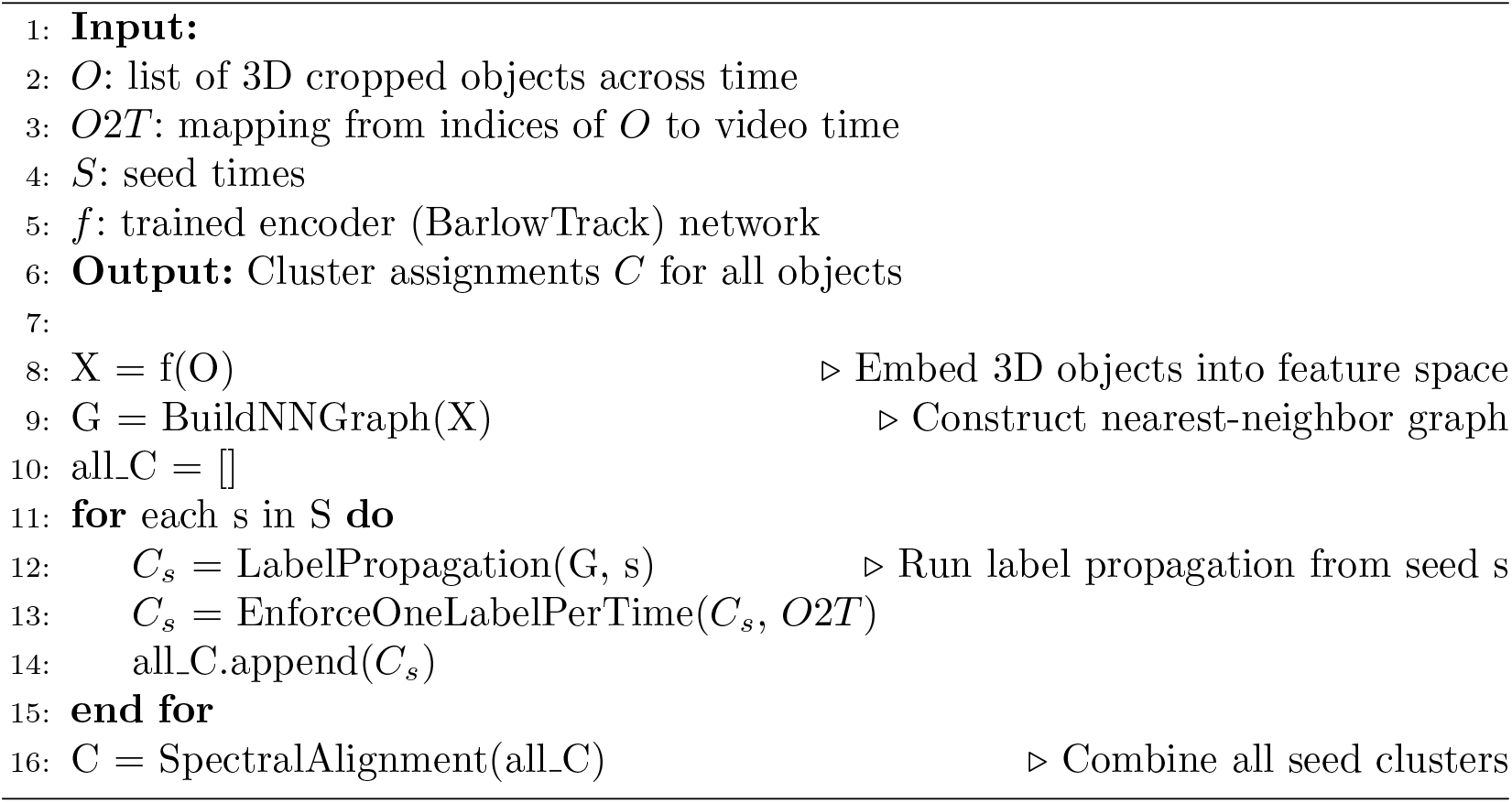

### 2.4 Results on benchmark datasets

Several scientific labs have published tracking algorithms^9,10,18–23^, and some have quantified the performance of these algorithms on ground truth. Published datasets with ground truth are summarized in Table 1, showing the variability of acquisition rates, number of volumes annotated, and spatial resolutions. Further, the images themselves are different (Fig. 3a-c). The Zimmer lab datasets^10^ have minimal preprocessing applied (only aligning images within a volume), and display significant body rotations across time (Figs. 1e, 3a). The single dataset from the Leifer lab^18^ displays a similar range of motion, but is recorded via a triangle wave scanner. This creates a small distortion in that the objects near the edges of the depth scan have a varying sampling rate, up to twice the sampling rate of objects in the center (which is equal to the average). The Flavell lab datasets^9^ have significant motion correction applied, and display a consistent body orientation (Fig. 3b). In both of these datasets, objects were segmented using a custom pipeline for the specific dataset; any subsequent tracking is limited by the performance of this segmentation. The large median inter-volume displacement in all cases makes traditional ROI-overlap based algorithms ineffective (Fig. S1a-b), and in each case the displacement is comparable to the nearest neighbor distance (Fig. S1c). Thus, an algorithm that directly learns features of new datasets is desirable.

**Fig. 1.**
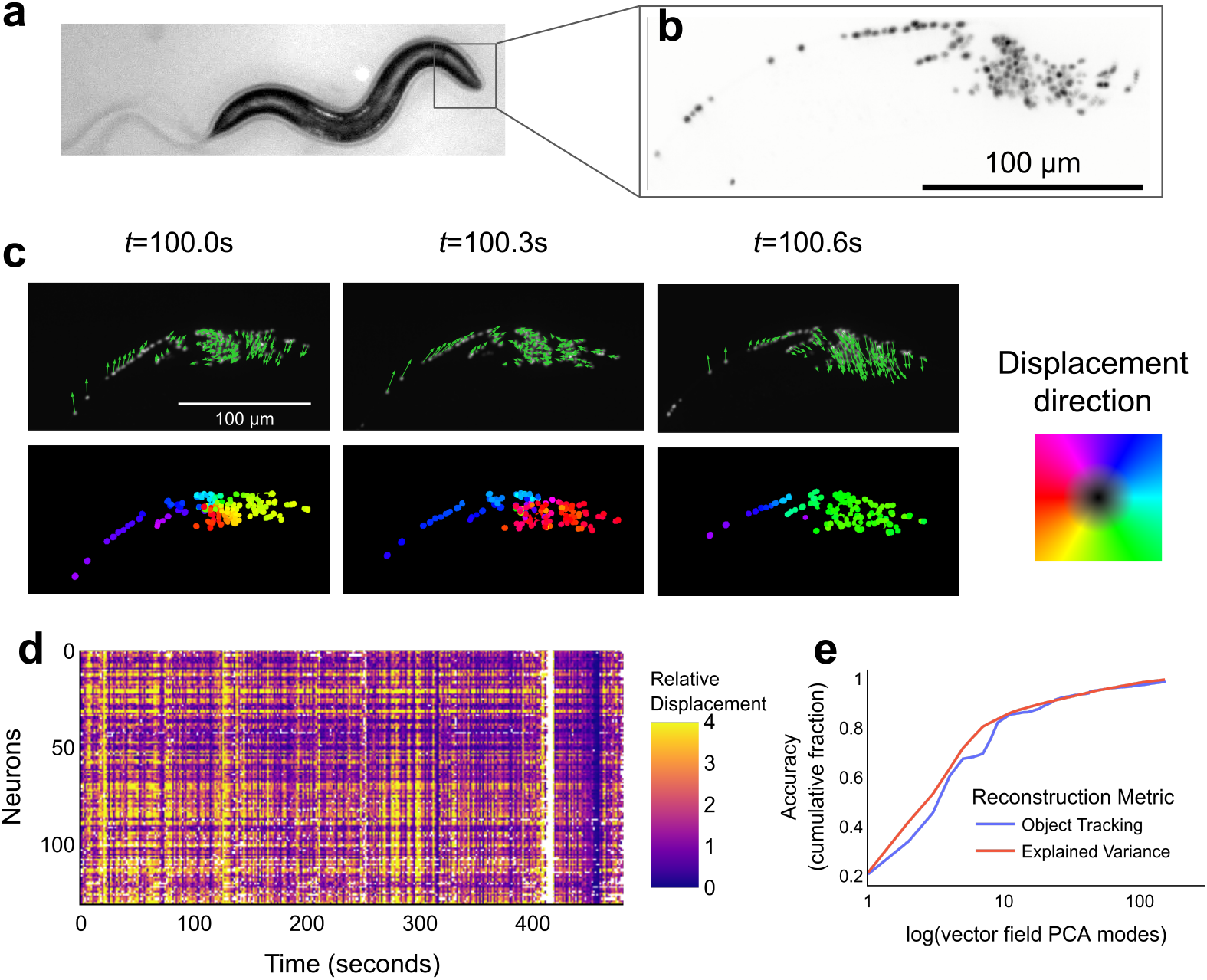
The tracking problem is complex and high dimensional. **(a)** Moving individual imaged using infrared illumination. **(b)** The head region is imaged with two channels at a high magnification, with raw data collected as 4D volumetric time series. For this work, only the reference channel is used. **(c)** Displacement vector fields and color-coded flow diagrams for 3 adjacent time points. **(d)** The displacement of individual neurons across time relative to the distance to their nearest neighbor. Large values are capped at 4, to better visualize variability. Gaps (white) showing time points missing in the ground truth. **(e)** Using Probabilistic PCA, the vector field across time of object displacements can be reconstructed; this measures the linearity (or linear dimensionality) of the motion. Shown are the variance explained and object tracking accuracy as a function of modes used for reconstruction.

**Fig. 2.**
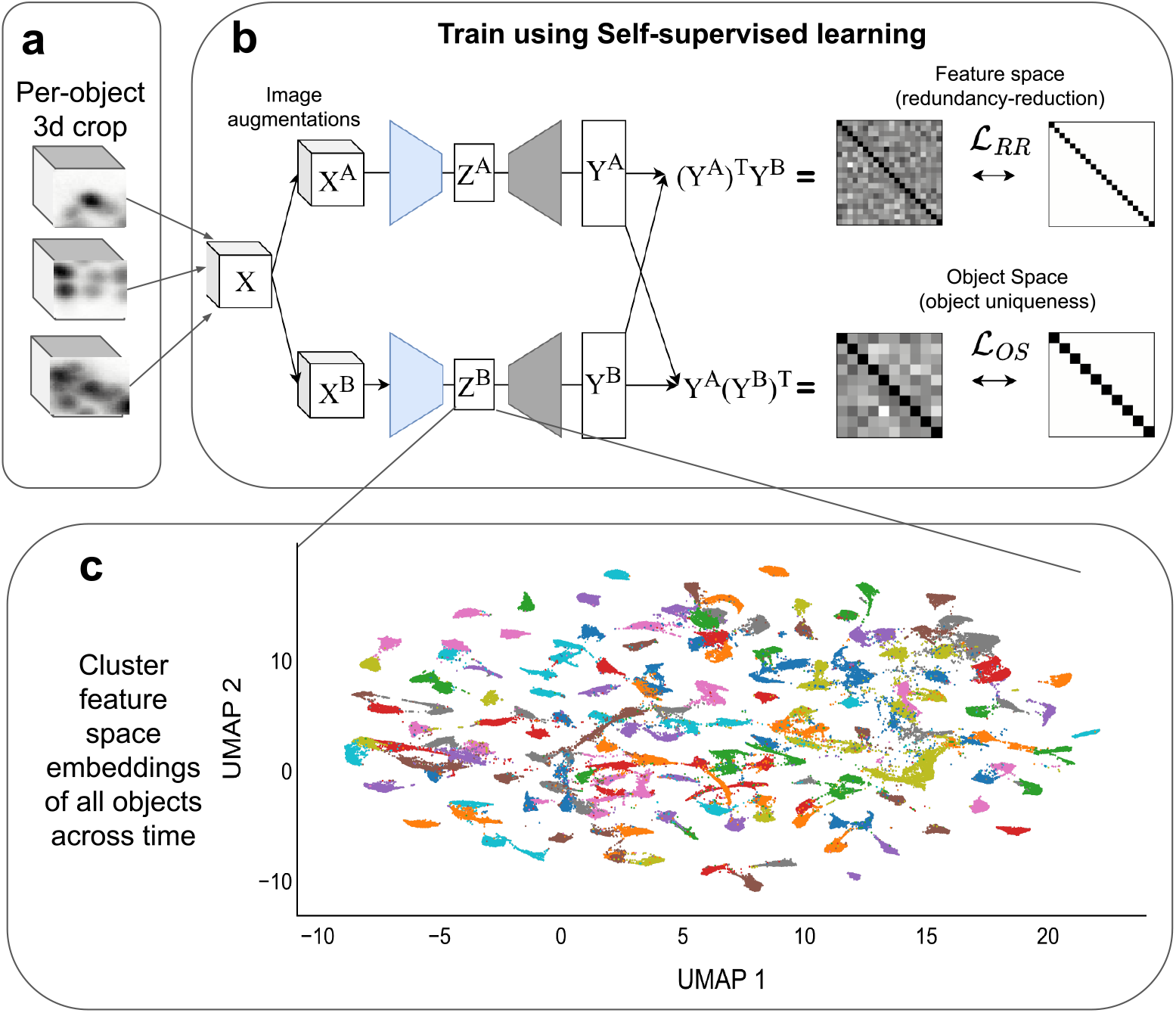
Algorithm design. **(a)** After object detection each object is cropped in 3d, including the neighboring region. **(b)** The neural network that embeds these crops into a latent space is trained using two loss terms: a redundancy reduction term calculated in feature space ( ℒ_*RR*_; same as [30]) as well as a novel loss term, calculated in object space (ℒ_*OS*_). *X*^*A*^ and *X*^*B*^ are two different image augmentations of the cropped neuron and neighborhood shown in (a). *Z* and *Y* are the feature space and projector space, respectively. The loss is applied to *Y*, but *Z* is used for downstream tasks. **(c)** Association of objects across time is achieved by clustering in *Z*, the latent feature space trained in (b). Colored by clustering (see methods); if successful, each cluster represents one object across the video. 6

**Fig. 3.**
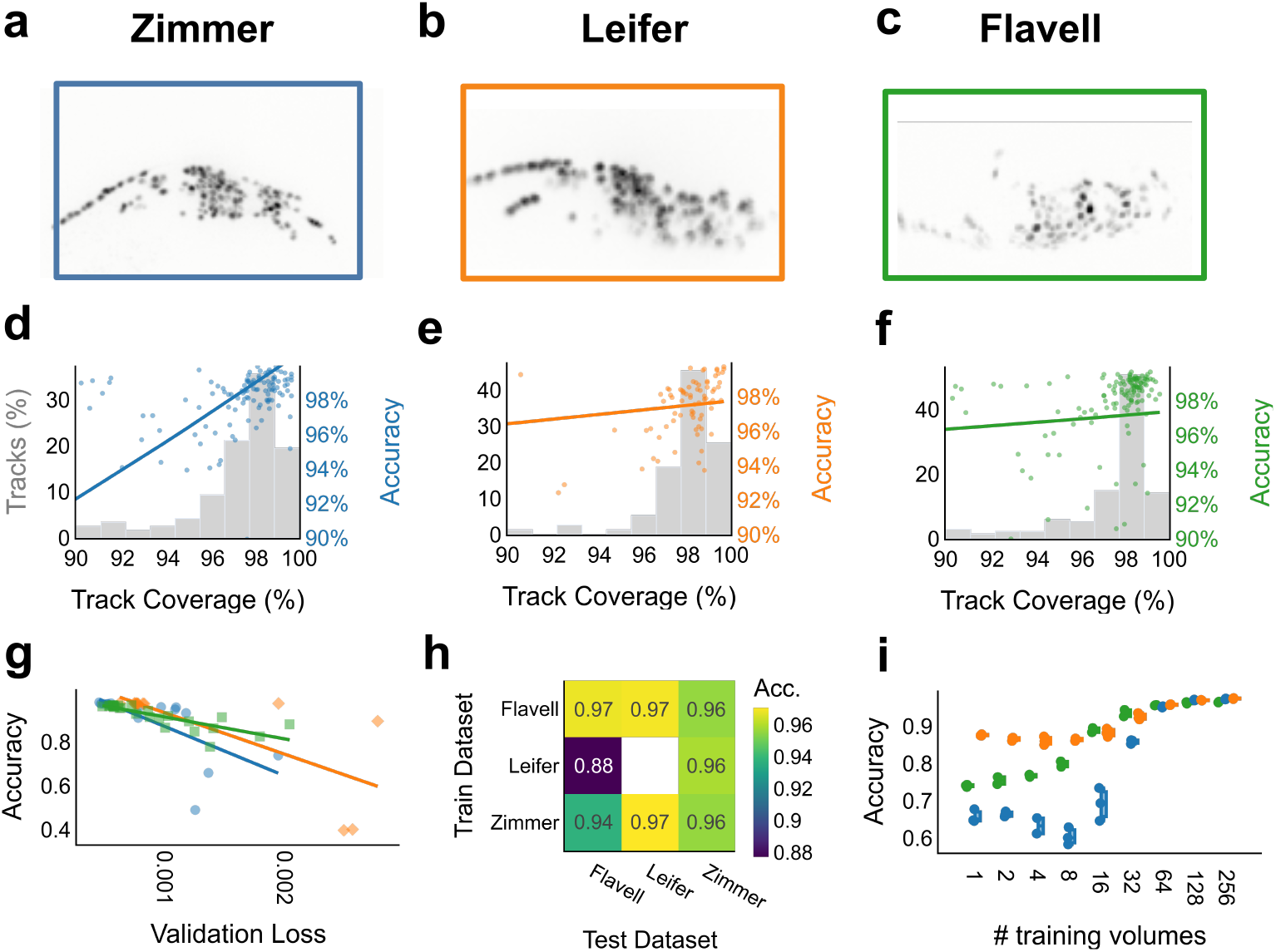
High performance and generalization on benchmark datasets. **(a-c)** Max z-projection of selected volumes for datasets from different scientific labs: (a) Zimmer, (b) Leifer, and (c) Flavell. **(d-f)** Quantifications of accuracy per track for the datasets in a-c, as a function of completeness of the track. Note that for the Flavell datasets, this is not manual annotation but rather their published algorithm[9]. **(g)** Negative relationship of validation loss and final tracking accuracy across a hyperparameter search. **(h)** Mean generalization accuracy of networks trained on one lab’s dataset to other datasets both within and between labs (a-c). There is only one dataset from the Leifer lab, and thus generalization accuracy cannot be calculated. **(i)** Accuracy as a function of number of training volumes used. Boxplots show median, interquartile range, and 1.5 times the interquartile range.

We applied our self-supervised learning algorithm, BarlowTrack, to each of these benchmark datasets. In each case we significantly improved the state of the art accuracy, without any manual annotation needed: greater than 96% (Table 2). We first sought to connect this accuracy to metrics that can be calculated on other datasets, for which no ground truth is present. For individual tracks, we found that the number of gaps is correlated to the accuracy (Fig. 3d-f, colored scatter plot), with our BarlowTrack pipeline achieved 98% coverage on most tracks (Fig. 3d-f, gray histogram). The test and validation losses from the network training itself provides another correlated proxy of the accuracy of the final network (Fig. 3g). Although there is no cost in human annotation to retrain a new network for every dataset, this is computationally expensive. Therefore we sought to quantify the generalization capabilities of a single trained network to datasets of the same conditions, and we found that similar accuracy was achieved by models trained on alternate datasets from the same lab (Fig. 3h). Generalization is also high across conditions, and for some users simply using a pretrained network may be sufficient. Together, users without access to expensive ground truth have both an alternative way to approximate their tracking accuracy, as well as confidence in the generalization capabilities of trained networks.

### 2.5 Ablation studies

We next sought to test the impact of various algorithmic choices. Using our custom clustering method gives a boost in accuracy compared to an off-the-shelf clustering method, and reduces the variance in accuracy of the resulting tracks (Fig. S1d). Both clustering methods tested are much better than the authors previous tracking method, which performs nearly at a noise level on the Flavell datasets (Fig. S1d). The accuracy was strongly dependent on the number of volumes used for training, with some datasets displaying a relatively sharp jump in performance around 64 volumes (around 4% of these videos) and saturating after 256 (around 16% of these videos; Fig. 3e). We further found that no single image augmentation was necessary, and removing any of them did not dramatically degrade performance (Fig. S1e).

Interestingly, using a completely untrained network and our proposed Label Propagation clustering method was sufficient to achieve a high level of performance, even similar to prior methods on some datasets (Fig. S1e). The Leifer dataset was the best tracked (86%), while the Zimmer dataset was not well tracked (62%). Including only a single augmentations highlighted the importance of Affine and Flip transformations, which generated networks with nearly full performance (Fig. S1f). In particular, the performance on the Leifer dataset did not degrade (97.4%) and only fell a few percentage on the Flavell data (to 94.2%). On the other hand, only using Blur, Noise, and Elastic Deformation were close in performance to completely untrained networks. Finally, we found that there was not a large difference between ablating either the original redundancy reduction or the novel loss term (Fig. S1e). Performing a hyperparameter sweep on the balance between these two showed that all accuracies were near the baseline, meaning that the sensitivity to this parameter was effectively 0. Thus instead of showing a large improvement via our novel loss term, we have shown that the original redundancy reduction loss term can be completely replaced without performance penalties.

All together, this algorithm achieves better than state-of-the-art performance across benchmark datasets, with a high degree of generalization and reusability.

### 2.6 Successful tracking in conditions without ground truth

With this algorithm in hand, we tried novel conditions that were previously inaccessible. Of particular interest are faster animals; In our previous work we deliberately created a higher density agarose pad to slow down the animals and make the tracking problem easier^10^; these density conditions were similar in other whole-brain imaging studies^9^. For this reason, we sought to track worms that move very quickly (Fig. 4a-c), which can be achieved by decreasing the density of the environment (reducing agarose concentration). Our faster condition displays more displacement per time point than the baseline condition. Another challenge is relatively simple changes in hardware settings, like increased magnification (Fig. 4d-f); this changes the motion between volumes (Fig. S2g-h) as well as the density of the objects in the image (Fig. S2i). A similar change in density in generated by using younger and smaller animals; all published studies thus far are of the young adult life stage of the animal. Larval life stage 3 (L3; young adult comes after L4) are only half the size of the young adult, making their brains more dense (Fig. S2i), but we could still achieve good performance (Fig. 4g-i). We further tried other interesting use cases, like a dimmer, more variable, and sparser reference channel promoter (the neuropal strain; Fig. S2a-c) and genetic strains with additional markers which changes the neighborhood features used for tracking the objects (Fig. S2d-f) In each case we achieved good performance (based on track coverage and overall heatmap quality), demonstrating the generalization capabilities of BarlowTrack.

**Fig. 4.**
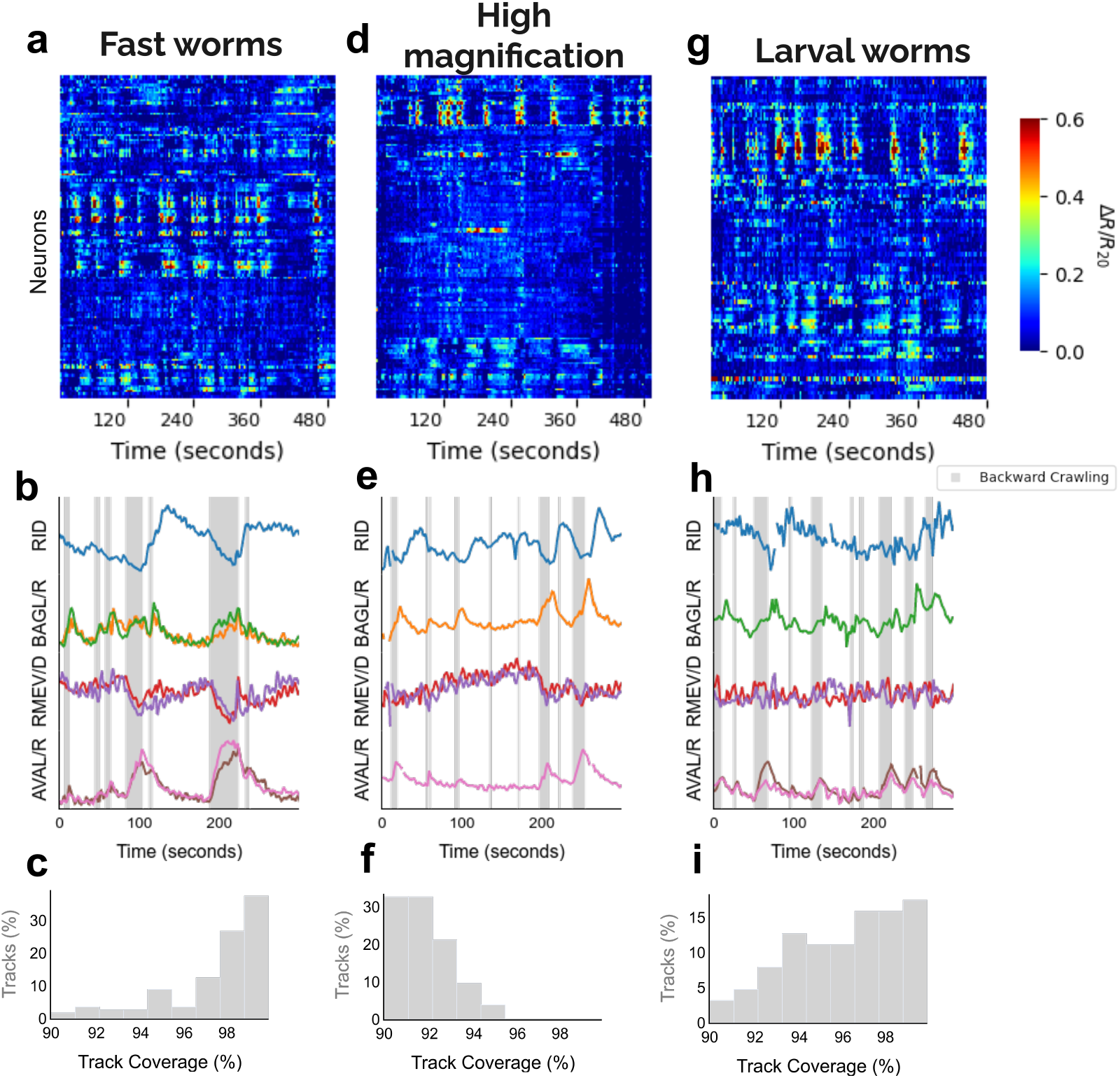
Performance on novel conditions. **(a)** Example heatmap of dataset in previously untracked conditions: an environmental change leading to a less viscous environment and fast animals (1**(b)** Traces of selected identifiable neurons for conditions in a). **(c)** Percent tracking coverage for individual tracked objects for datasets in a), analogous to Fig. 3d-f but without ground truth **(d-f)** Same as (a-c), but for a different condition: increased magnification (63x vs. 40x). **(g-i)** Same as (a-c), but for a different condition: a younger life stage of the worm, larval stage 3 (L3).

For all of these cases, segmentation or object detection limits the performance of the BarlowTrack network, both during training and tracking. However, for this study a single StarDist network trained on 10 partially segmented volumes of Zimmer datasets was sufficient to achieve good performance. Such high quality tracking of novel conditions allows rapid testing and exploration of new hypotheses.

### 2.7 GUI-based evaluation and correction

To facilitate interpretation, we provide a Napari-based^48^ graphical user interface compatible with the NWB format (Fig. 5a). This allows visualization of the different layers produced by or required for the tracking pipeline, including multi-channel raw data, raw and tracked segmentation (Fig. 5b), tracking information, and neuron IDs ((Fig. 5c).

**Fig. 5.**
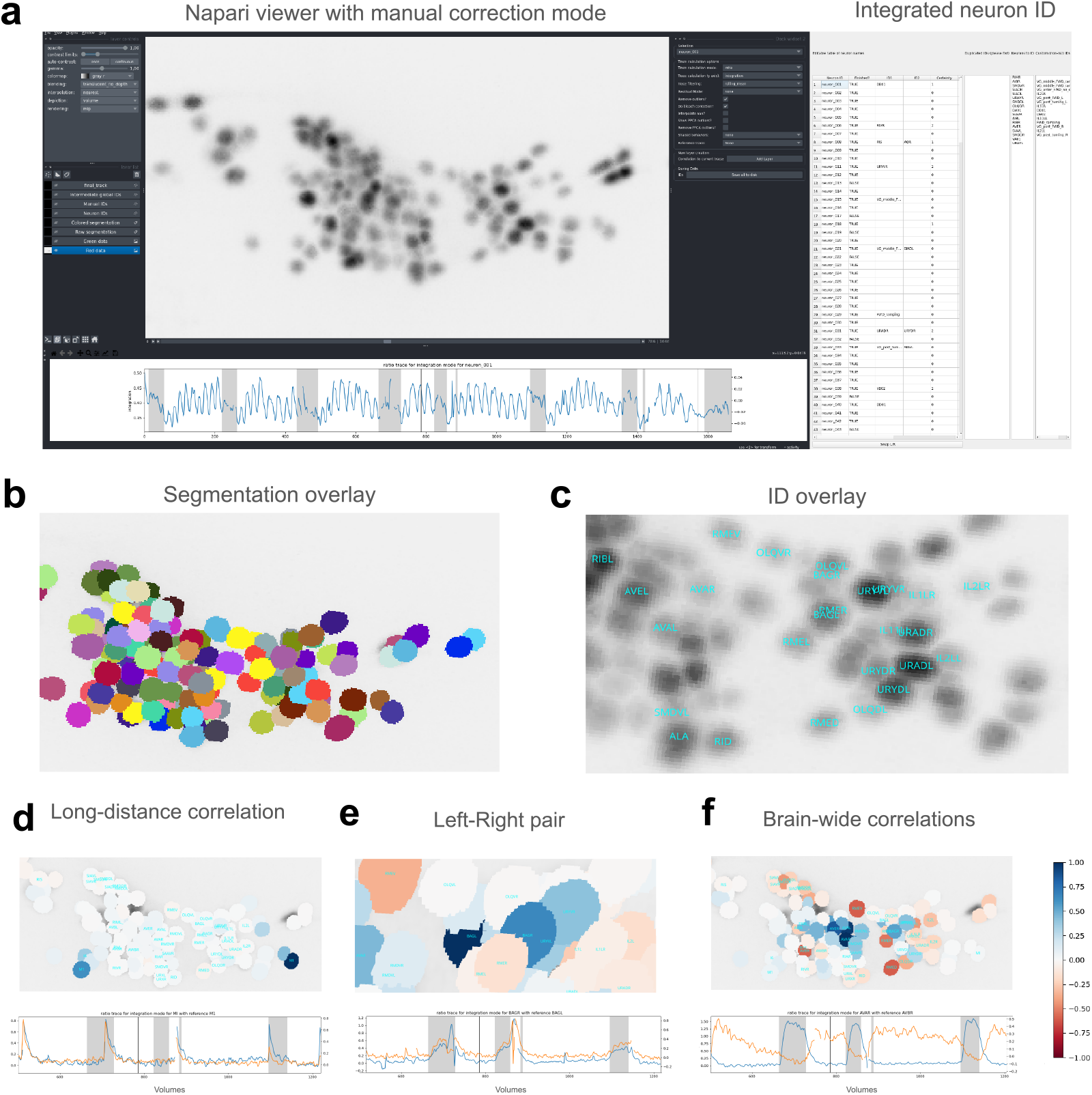
Overview of the GUI. **(a)** Napari-based GUI with multiple image layers (left), trace of a selected neuron (bottom), and a dedicated panel for IDing neurons (right). **(b)** Example view of the segmentation overlay. **(c)** Example view of the neuron ID overlay, which can be modified live via the IDing panel shown in (a). **(d-f)** Neuron time series activity overlay with specific workflows, where blue identifies correlated activity and red anti-correlated activity. (d) Separated but highly correlated neurons. (e) A Left-Right pair of neurons. (f) A widely distributed, correlated signal. For each case, two neuron traces are shown in the GUI, corresponding to the dark blue neuron for the blue trace (100% correlated with itself).

Identification of neurons using globally expressed genetic markers like Neuropal^49^ is powerful and has been used in many cases to great success. However, for some experiments it is not feasible to use this strain (e.g. locomotion deficit of the animal^36,50^ or lack of wild-type sensory activity^10^), and other identification workflows are required. We provide a tool to support position and time-series based annotation, specifically using the correlation between an anchor neuron and all others, shown via recoloring the segmentation at one time point. Using this view, very strong correlations can be immediately seen, and, depending on position, the neurons can be identified. For example, several pharyngeal neurons have unique positions on opposite sides of the brain, but are highly correlated (Fig. 5d). Left-Right neuron pairs are often easy to identify (Fig. 5e) via activity. Because the animal is on it’s ventral or dorsal side, these pairs are on the top or bottom of the image; such pairs usually have extremely correlated activity (usually *>* 0.95^51^). However, deformations can lead to offset or distinct positions, and thus the correlation information supplements the position information to allow confident identification. As a much more complex example, some signals are widely distributed among both neighboring and spatially separated neurons (Fig. 5f). However, it is important to be cautious with this method of identification. Especially when neurons with correlated activity occur in the same regions, single-neuron^52^ or Neuropal^9^ studies must be done to confirm the activity of the neuron, and relative positioning must be confirmed to ensure neighboring neurons do not stochastically switch position^31,49^, as is known for several regions. With these points of caution, a set of neurons can be identified without genetic markers; see prior work^10,51,52^ for more details. For trained users, of set of 60 neurons commonly identified in the Zimmer lab^10^ can be completed in 15 minutes, which previously took several hours.

Tracking mistakes can be corrected using an alternative tracklet-based approach; although the method may not generalize to all conditions (Fig. S1d) it is much easier to manually correct. In our tests^10^ this method of generating ground truth is 10-100x faster than per-timepoint correction used in other studies^20,21,24^ In short, tracklet-based tracking generates high quality but short time “tracklets” which can be checked more quickly; see our prior publication for more details^10^. The full track is then generated by associating tracklets at different time points, and thus the main task of manual annotation is to check that this association is correct and occasionally to split tracklets apart. We have produced several YouTube videos detailing this workflow (See tutorials and text on GitHub^53^).

Taken together, we provide a tracking method and analysis GUI that allows exploration of never-before-seen datasets, including new life stages of *C. elegans* and novel laboratory and experimental conditions.

## 3 Discussion

Microscopy has advanced such that it is now possible to collect data from nearly every neuron in the body of small, transparent, freely moving organisms such as *C. elegans*. However, the software to reliably analyze this data has been limited to the highly specific conditions of single labs and, even within labs, specific experimental and hardware conditions. We present an algorithm that uses self-supervised learning to learn a feature space directly from data, solving the generalization problem.

The success of this approach is based in the local similarity of deformations undergone by the brain and the augmentations used in the training procedure. Thus, the algorithm is not specific to *C. elegans*, and could be used to track other species.

However, if a brain undergoes very different deformations, such as the dramatic expansion and contraction of a hydra brain^8^, this approach may not perform well without additional augmentations. If individual objects are distinct enough that neighborhood information is not needed, then they can still be successfully clustered. Thus, as datasets become available, this method could be immediately applied to other similar species such as *C. briggsae* and *Pristionchus pacificus*, as well as significantly different organisms with a similar body pattern like *Ciona* or *Tardigrades*^54^. Further, this approach generates a feature space which can be used as a high quality starting point for other tracking algorithms.

A major bottleneck of all detect-then-track approaches is object detection or, equivalently for this case, instance segmentation. Although there is significant work on generalization in that field^16,17^, the densest brain regions in *C. elegans* cannot be reliably segmented with current methods due to complete merging of the regions of interest. This leads to a loss of 10-40% of the neurons in the head, depending on imaging resolution and size of the animal. A promising area of future work is to use self-supervised learning for this problem as well^55^ or to leverage video data more directly via, for example, the Mask RCNN architecture^56^.

This workflow uses a CNN architecture, and thus generates a feature space purely from image information. A clear avenue of future work is to include position information, for example using established transformer architectures^57^. Such approaches have already been applied to this tracking problem and shown promising results^10,22^. Using such information, this algorithm may generalize to even more complex situations, such as social studies with the presence of multiple animals^24^.

We have shown that this method is broadly applicable, re-released published benchmark datasets in a standardized NWB format, and provided visualization software to interactively view the videos and traces. Our approach will contribute to a democratization and standardization of whole-brain freely moving experiments.

## 4 Methods

### 4.1 Explanation of datasets

See Table 1 for an overview of the datasets, and the referenced papers for more information regarding preprocessing performed on the images. A short summary of the relevant differences is given below.

The Zimmer lab datasets use an evenly expressed promoter, leading to bright and uniform expression in the reference channel^10^. This allows high-quality segmentation via published methods (StarDist^12^), but leads to more similarity between different local neighborhoods of specific neurons. The only image preprocessing steps applied were background subtraction and rigid alignment between z-slices, which partially corrected for intra-volume motion. Z scanning was performed with a sawtooth wave, leading to low-quality “flyback frames” which were removed from the final video; speecifically, out of 24 z-slices recorded, 2 were dropped. Semi-automatic ground truth was generated using the GUI presented in Fig. 5, with more details in prior work^10^.

Similarly, the Leifer lab datasets are not significantly preprocessed. Z scanning was performed using a triangle wave, but the video was reindexed to produce a consistent z depth in that dimension. This means that in alternating volumes, the exact timing of individual slices is different; this is neglected in the tracking problem. Ground truth was generated via semi-automated annotation, with more details in prior work^18^.

The Flavell lab datasets use a different pan-neuronal promoter, which is heterogeneously expressed^9^. This leads to more unique neighborhoods around each neuron, but some neurons are nearly invisible. A custom Unet-based segmentation algorithm was needed to produce good results. However, it sometimes oversegmented the neurons; this was corrected for the custom tracking algorithm of the Flavell lab, but is not supported by this work. This means that neurons corresponding to multiple segmented objects were dropped from the ground truth. Significant nonrigid motion correction was applied, which stabilized the heading of the animal to be the same for the majority of the video. Rare 180-degree flips occur, completely changing the neighborhood features of neurons, leading to the particular importance of 180-flip augmentation in training. No ground truth was generated for these datasets, but rather a stochastically expressed GFP (non-time-varying) was used to benchmark tracking mismatches.

### 4.2 Calculation of object reconstruction accuracy

When reconstructing the vector field of the motion from linear modes, variance explained may not be sufficient to quantify tracking performance. This is because objects in denser regions must be more accurately reconstructed to prevent identity swaps. Thus, the object reconstruction accuracy is calculated as the fraction of objects whose reconstruction error is less than half the nearest neighbor distance, calculated per time point.

### 4.3 Hyperparameter tuning

#### 4.3.1 BarlowTrack training

We performed hyperparameter sweeps for each dataset varying the following parameters, with the chosen values shown:

- learning rate: 0.0001
- projector final (the dimensionality of the final projector output layer, on which the loss is calculated): 512
- embedding dim (the dimensionality of the internal feature space, used for later
- clustering): 64
- target_sz_z (the number of z slices used in crops): 8
- target_sz_xy (the size in xy of the slices used in crops): 64
- *λ_obj* (the balance between feature- and object-space loss): 0.67

We did not find a strong dependence on final accuracy for the dimensionality, loss balance, or learning rate parameters; more discussion of the embedding dimension is in the next section. The size of the crop however was very important, with small crops not performing well. Further, due to the increased z resolution of the Flavell datasets, a proportional increase in z slices was key for good performance (16 instead of 8).

#### 4.3.2 Clustering

Clustering algorithms generally need to operate on a low dimensional space to be efficient. We therefore set the dimension of the learned embedding space to 64; lower than this generated worse performance in hyperparameter testing. For HDBSCAN, the space is further reduced in dimension linearly via PCA (to 50) and then non-linearly via UMAP^58^ (to 10).

We further used the following parameters for these algorithms:

- HDBSCAN (step 1: UMAP): n_neighbors=10, min_dist=0
- HDBSCAN (step 2: clustering): min_cluster_size=0.5*N, min_samples=0.02*N, cluster_selection_method=‘leaf’
- Label Propagation (step 1: graph formation): n_neighbors=20
- Label Propagation (step 2: label propagation): n_layers=100, top k=2, num_seeds=100
- Label Propagation (step 3: label alignment): confidence_threshold=0.1

where N is the total number of frames in the video, n layers is the number of label propagation steps to take, and top k is the number of candidate labelings to keep for the next step. Other parameters are directly passed to the relevant libraries.

### 4.4 Calculation of time series

Unless otherwise stated, time series data were calculated as the sum of all pixels in the signal channel divided by the sum of all pixels in the reference channel, after background subtraction. These ratiometric time series were then bleach corrected. This is similar to the prior published time series for the benchmark datasets. The Leifer lab datasets^18^ did not include segmentation, and traces were not recalculated for them.

**Supplementary information**.

### 4.5 Data and Code Availability

All original code for video processing is available at: https://github.com/Zimmer-lab/barlow track^53^ Data will be made available upon publication.

## 4.6 Acknowledgements

The computational results of this work have been achieved using the Life Science Compute Cluster (LiSC) of the University of Vienna. The research leading to these results has received funding from the European Research Council (ERC) under the European Union’s Horizon 2020 research and innovation programme (M.Z. and J.M. *elegans*BrainBodyEnvi, #101054527), from the Simons Foundation (#543069 and NC-GB-CULM-00003196-01 for M.Z., AN-SURFiN-00008221 and AN-SURFiN-00015263 for Z.Z.), the University of Vienna, and the Research Institute of Molecular Pathology (IMP). C.F. and I.L. were supported by VIP2 postdoctoral fellowships co-funded by the European Union’s Horizon 2020 research and innovation programme under the Marie Sklodowska-Curie grant agreement No. 847548. I.L. thanks the support from the EMBO funding agency (ALTF 1037-2019) and the Human Frontier Science Program fellowship (LT000335/2020-L).

## 4.7 Author Contributions

C.F. conceived the computational methods and design, and wrote the manuscript. C.F. and S.S developed the computational methods. I.L., J.M., and Z.Z. performed experiments. M.Z. led the studies.

**Fig. S1.**
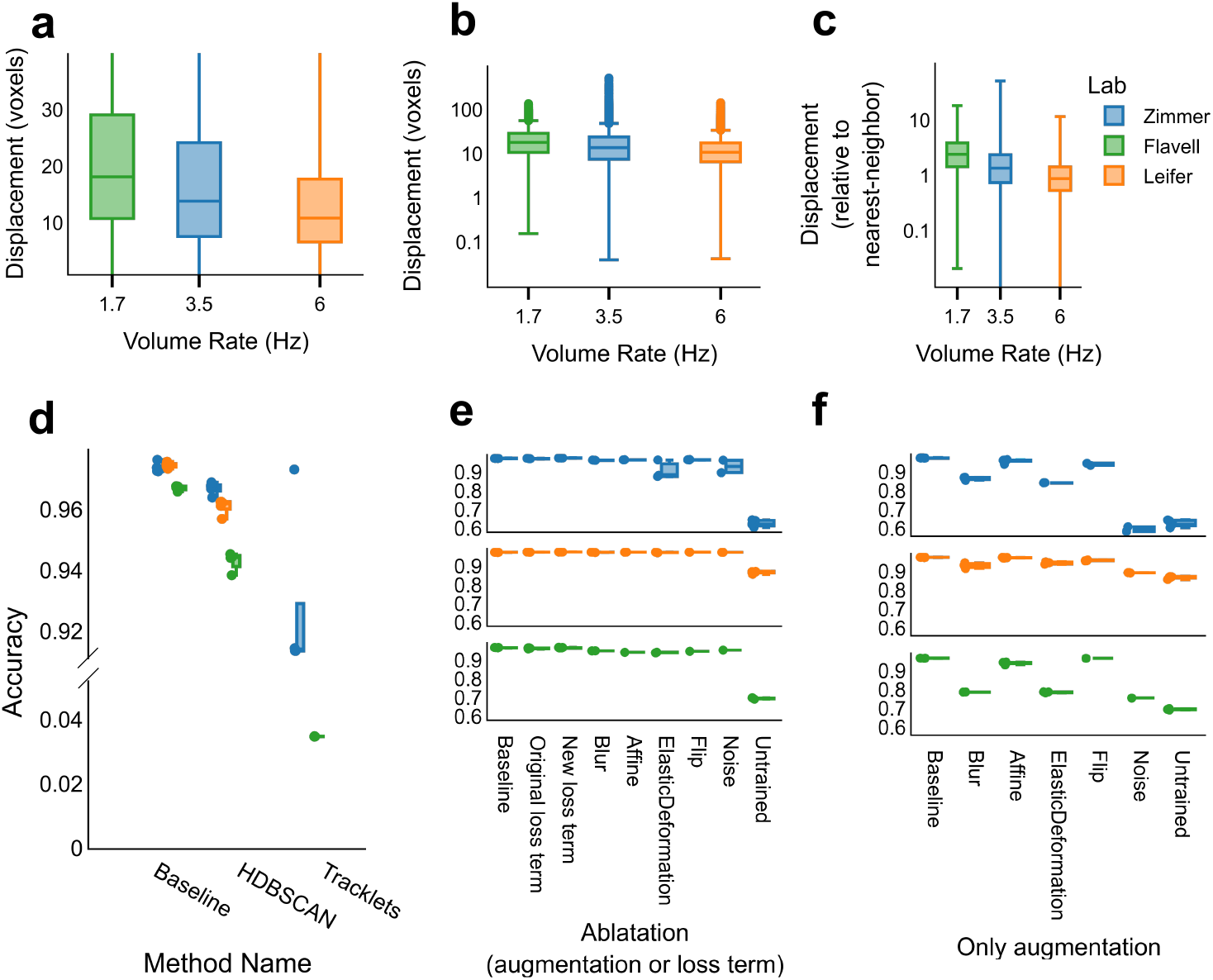
Further quantification of performance on benchmark datasets. **(a-c)** Inter-volume displacement of neurons in the ground truth (for the Flavell datasets, this is not manual annotation but rather their published algorithm[9]). Panels (a-b) are the same information, but a) has the outliers removed in order to show the different medians more clearly. b) shows the full range. Panel c) is normalized by the nearest neighbor distance. **(d)** Comparison between pipelines. HDBSCAN clustering uses the same BarlowTrack network as the Baseline; the only difference is the clustering algorithm (Baseline uses the Label Propagation method described the main text). The tracklet method is from the authors prior work[10], and was trained on the Zimmer datasets used for evaluation. Note the near useless performance (broken y-axis) on the Flavell dataset. **(e-f)** Baseline network and ablation studies of different training data augmentation, including the novel loss term described in Fig. 2b. e) shows ablation of specific augmentations or loss terms, while f) show single additions. All panels with boxplots show median, interquartile range, and 1.5 times the interquartile range.

**Fig. S2.**
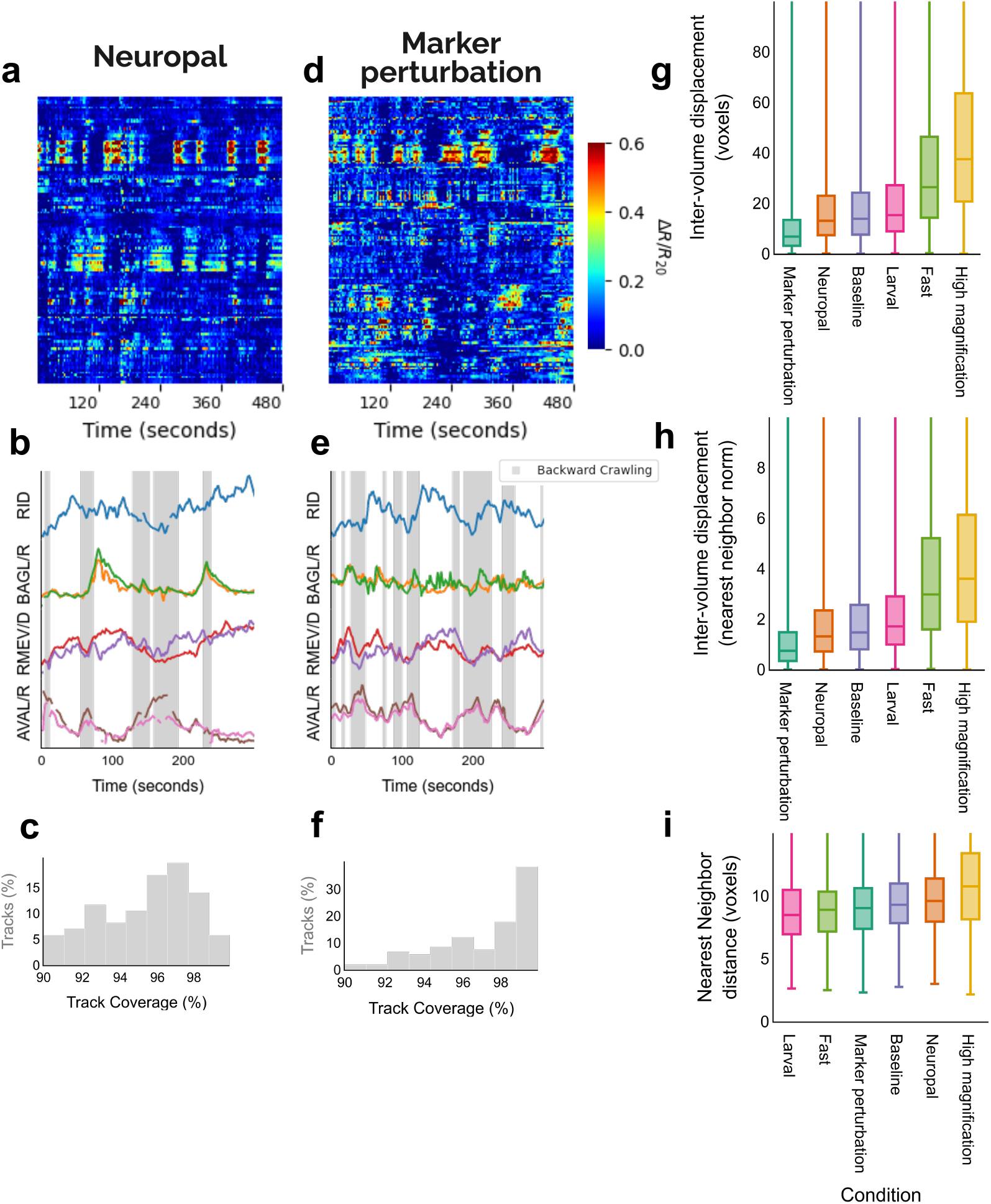
Performance on additional novel conditions. **(a-c)** Example heatmaps, traces, and tracking quality of previously difficult-to-track condition: the neuropal genetic background, with variable and dim reference markers. This is the same genetic background as the Flavell datasets. **(d-f)** Same as (a-c) but with an additional genetic marker, changing the neighborhood of many neurons. **(g)** Intervolume displacement of the different conditions. **(h)** Same as g) but with displacement normalized to nearest neighbor distance. **(i)** The nearest-neighbor distance for each condition. All panels with boxplots show median, interquartile range, and 1.5 times the interquartile range.

## References

1 Churchland, M. M., J. P. Cunningham, M. T. Kaufman, J. D. Foster, P. Nuyujukian, S. I. Ryu, and K. V. Shenoy (2012). “Neural population dynamics during reaching”. In: Nature 487.7405, pp. 51–56.

2 Aimon, S., K. Y. Cheng, J. Gjorgjieva, and I. C. G. Kadow (2023). “Global change in brain state during spontaneous and forced walk in Drosophila is composed of combined activity patterns of different neuron classes”. In: Elife 12, e85202.

3 Brezovec, B. E., A. B. Berger, Y. A. Hao, F. Chen, S. Druckmann, and T. R. Clandinin (2024). “Mapping the neural dynamics of locomotion across the drosophila brain”. In: Current Biology 34.4, pp. 710–726.

4 Steinmetz, N. A., P. Zatka-Haas, M. Carandini, and K. D. Harris (2019). “Distributed coding of choice, action and engagement across the mouse brain”. In: Nature 576.7786, pp. 266–273.

5 Zong, W., H. A. Obenhaus, E. R. Skytøen, H. Eneqvist, N. L. de Jong, R. Vale, M. R. Jorge, M.-B. Moser, and E. I. Moser (2022). “Large-scale two-photon calcium imaging in freely moving mice”. In: Cell 185.7, pp. 1240–1256.

6 Skocek, O., T. Nöbauer, L. Weilguny, F. Martínez Traub, C. N. Xia, M. I. Molodtsov, A. Grama, M. Yamagata, D. Aharoni, D. D. Cox, et al. (2018). “High-speed volumetric imaging of neuronal activity in freely moving rodents”. In: Nature methods 15.6, pp. 429–432.

7 Kim, D. H., J. Kim, J. C. Marques, A. Grama, D. G. Hildebrand, W. Gu, J. M. Li, and D. N. Robson (2017). “Pan-neuronal calcium imaging with cellular resolution in freely swimming zebrafish”. In: Nature methods 14.11, pp. 1107–1114.

8 Lagache, T., A. Hanson, J.E. Pérez-Ortega, A. Fairhall, and R. Yuste (2021). “Tracking calcium dynamics from individual neurons in behaving animals”. In: PLoS computational biology 17.10, e1009432.

9 Atanas, A. A., J. Kim, Z. Wang, E. Bueno, M. Becker, D. Kang, J. Park, T. S. Kramer, F. K. Wan, S. Baskoylu, et al. (2023). “Brain-wide representations of behavior spanning multiple timescales and states in C. elegans”. In: Cell 186.19, pp. 4134–4151.

10 Fieseler, C., I. Lev, U. Rey, L. Hille, H. Brenner, and M. Zimmer (2025). “An intrinsic neuronal manifold underlies brain-wide hierarchical organization of behavior in C. elegans”. In: bioRxiv, pp. 2025–03.

11 Hallinen, K. M., R. Dempsey, M. Scholz, X. Yu, A. Linder, F. Randi, A. K. Sharma, J. W. Shaevitz, and A. M. Leifer (July 2021). “Decoding locomotion from population neural activity in moving C. elegans”. In: eLife 10. Ed. by R.L. Calabrese. Publisher: eLife Sciences Publications, Ltd, e66135. issn: 2050-084X. doi: 10.7554/eLife.66135.

12 StarDist Object Detection with Star-convex Shapes (Nov. 2022). original-date: 2018-06-26T19:54:45Z.

13 Weigert, M., U. Schmidt, R. Haase, K. Sugawara, and G. Myers (Mar. 2020). “Star-convex Polyhedra for 3D Object Detection and Segmentation in Microscopy”. en. In: 2020 IEEE Winter Conference on Applications of Computer Vision (WACV). Snowmass Village, CO, USA: IEEE, pp. 3655–3662. isbn: 978-1-72816-553-0. doi: 10.1109/WACV45572.2020.9093435.

14 Schmidt, U., M. Weigert, C. Broaddus, and G. Myers (2018). “Cell detection with star-convex polygons”. In: International Conference on Medical Image Computing and Computer-Assisted Intervention. Springer, pp. 265–273.

15 Stringer, C., T. Wang, M. Michaelos, and M. Pachitariu (2021). “Cellpose: a generalist algorithm for cellular segmentation”. In: Nature methods 18.1, pp. 100– 106.

16 Pachitariu, M., M. Rariden, and C. Stringer (2025). “Cellpose-SAM: superhuman generalization for cellular segmentation”. In: bioRxiv, pp. 2025–04.

17 Archit, A., L. Freckmann, S. Nair, N. Khalid, P. Hilt, V. Rajashekar, M. Freitag, C. Teuber, M. Spitzner, C. Tapia Contreras, et al. (2025). “Segment anything for microscopy”. In: Nature Methods 22.3, pp. 579–591.

18 Nguyen, J. P., F. B. Shipley, A. N. Linder, G. S. Plummer, M. Liu, S. U. Setru, J. W. Shaevitz, and A. M. Leifer (2016). “Whole-brain calcium imaging with cellular resolution in freely behaving Caenorhabditis elegans”. In: Proceedings of the National Academy of Sciences 113.8, E1074–E1081.

19 Wu, Y., S. Wu, X. Wang, C. Lang, Q. Zhang, Q. Wen, and T. Xu (2022). “Rapid detection and recognition of whole brain activity in a freely behaving Caenorhabditis elegans”. In: PLoS computational biology 18.10, e1010594.

20 Ryu, J., A. Nejatbakhsh, M. Torkashvand, S. Gangadharan, M. Seyedolmohadesin, J. Kim, L. Paninski, and V. Venkatachalam (2024). “Versatile multiple object tracking in sparse 2D/3D videos via deformable image registration”. In: PLOS Computational Biology 20.5, e1012075.

21 Park, C. F., M. Barzegar-Keshteli, K. Korchagina, A. Delrocq, V. Susoy, C. L. Jones, A. D. Samuel, and S. J. Rahi (2024). “Automated neuron tracking inside moving and deforming C. elegans using deep learning and targeted augmentation”. In: Nature Methods 21.1, pp. 142–149.

22 Yu, X., M. S. Creamer, F. Randi, A. K. Sharma, S. W. Linderman, and A. M. Leifer (2021). “Fast deep neural correspondence for tracking and identifying neurons in C. elegans using semi-synthetic training”. In: Elife 10, e66410.

23 Atanas, A. A., A. K.-Y. Lu, B. Goodell, J. Kim, S. Baskoylu, D. Kang, T. S. Kramer, E. Bueno, F. K. Wan, K. L. Cunningham, et al. (2024). “Deep Neural Networks to Register and Annotate Cells in Moving and Deforming Nervous Systems”. In: bioRxiv, pp. 2024–07.

24 Susoy, V., W. Hung, D. Witvliet, J. E. Whitener, M. Wu, C. F. Park, B. J. Graham, M. Zhen, V. Venkatachalam, and A. D. Samuel (2021). “Natural sensory context drives diverse brain-wide activity during C. elegans mating”. In: Cell 184.20, pp. 5122–5137.

25 Nejatbakhsh, A., E. Varol, E. Yemini, V. Venkatachalam, A. Lin, A. D. Samuel, and L. Paninski (2020). “Extracting neural signals from semi-immobilized animals with deformable non-negative matrix factorization”. In: bioRxiv, pp. 2020–07.

26 Skuhersky, M., T. Wu, E. Yemini, A. Nejatbakhsh, E. Boyden, and M. Tegmark (2022). “Toward a more accurate 3D atlas of C. elegans neurons”. In: BMC bioinformatics 23.1, p. 195.

27 Wen, C., T. Miura, V. Voleti, K. Yamaguchi, M. Tsutsumi, K. Yamamoto, K. Otomo, Y. Fujie, T. Teramoto, T. Ishihara, et al. (2021). “3DeeCellTracker, a deep learning-based pipeline for segmenting and tracking cells in 3D time lapse images”. In: Elife 10, e59187.

28 Maška, M., V. Ulman, P. Delgado-Rodriguez, E. Gómez-de-Mariscal, T. Nečasová, F. A. Guerrero Peña, T. I. Ren, E. M. Meyerowitz, T. Scherr, K. Löffler, et al. (2023). “The cell tracking challenge: 10 years of objective benchmarking”. In: Nature Methods 20.7, pp. 1010–1020.

29 Jaiswal, A., A. R. Babu, M. Z. Zadeh, D. Banerjee, and F. Makedon (2020). “A survey on contrastive self-supervised learning”. In: Technologies 9.1, p. 2.

30 Zbontar, J., L. Jing, I. Misra, Y. LeCun, and S. Deny (2021). “Barlow twins: Self-supervised learning via redundancy reduction”. In: International conference on machine learning. PMLR, pp. 12310–12320.

31 Sprague, D. Y., K. Rusch, R. L. Dunn, J. M. Borchardt, S. Ban, G. Bubnis, G. C. Chiu, C. Wen, R. Suzuki, S. Chaudhary, et al. (2025). “Unifying community whole-brain imaging datasets enables robust neuron identification and reveals determinants of neuron position in C. elegans”. In: Cell Reports Methods.

32 Stephens, G. J., B. Johnson-Kerner, W. Bialek, and W. S. Ryu (2008). “Dimensionality and dynamics in the behavior of C. elegans”. In: PLoS computational biology 4.4, e1000028.

33 Ilin, A. and T. Raiko (2010). “Practical approaches to principal component analysis in the presence of missing values”. In: The Journal of Machine Learning Research 11, pp. 1957–2000.

34 Voleti, V., K. B. Patel, W. Li, C. Perez Campos, S. Bharadwaj, H. Yu, C. Ford, M. J. Casper, R. W. Yan, W. Liang, et al. (2019). “Real-time volumetric microscopy of in vivo dynamics and large-scale samples with SCAPE 2.0”. In: Nature methods 16.10, pp. 1054–1062.

35 Bouchard, M. B., V. Voleti, C. S. Mendes, C. Lacefield, W. B. Grueber, R. S. Mann, R. M. Bruno, and E. M. Hillman (2015). “Swept confocally-aligned planar excitation (SCAPE) microscopy for high-speed volumetric imaging of behaving organisms”. In: Nature photonics 9.2, pp. 113–119.

36 Randi, F., A. K. Sharma, S. Dvali, and A. M. Leifer (2023). “Neural signal propagation atlas of Caenorhabditis elegans”. In: Nature 623.7986, pp. 406–414.

37 Karbowski, J., C. J. Cronin, A. Seah, J. E. Mendel, D. Cleary, and P. W. Sternberg (2006). “Conservation rules, their breakdown, and optimality in Caenorhabditis sinusoidal locomotion”. In: Journal of theoretical biology 242.3, pp. 652–669.

38 Sawin, E. R., R. Ranganathan, and H. R. Horvitz (2000). “C. elegans locomotory rate is modulated by the environment through a dopaminergic pathway and by experience through a serotonergic pathway”. In: Neuron 26.3, pp. 619–631.

39 Ben Arous, J., S. Laffont, and D. Chatenay (2009). “Molecular and sensory basis of a food related two-state behavior in C. elegans”. In: PloS one 4.10, e7584.

40 Rübel, O., A. Tritt, R. Ly, B. K. Dichter, S. Ghosh, L. Niu, P. Baker, I. Soltesz, L. Ng, K. Svoboda, et al. (2022). “The Neurodata Without Borders ecosystem for neurophysiological data science”. In: Elife 11, e78362.

41 Adhinarta, J., J. Dong, T. He, J. Huang, D. Sprague, J. Wan, H. J. Lee, Z. Yu, H. Lu, E. Yemini, et al. (2025). “WormID-Benchmark: Extracting Whole-Brain Neural Dynamics of C. elegans At the Neuron Resolution”. In: bioRxiv, pp. 2025– 01.

42 McInnes, L., J. Healy, and S. Astels (Mar. 2017). “hdbscan: Hierarchical density based clustering”. In: The Journal of Open Source Software 2.11. doi: 10.21105/joss.00205.

43 Wolny, A., L. Cerrone, A. Vijayan, R. Tofanelli, A. V. Barro, M. Louveaux, C. Wenzl, S. Strauss, D. Wilson-Sánchez, R. Lymbouridou, et al. (July 2020). “Accurate and versatile 3D segmentation of plant tissues at cellular resolution”. In: eLife 9. Ed. by C. S. Hardtke, D. C. Bergmann, D. C. Bergmann, and M. Graeff, e57613. issn: 2050-084X. doi: 10.7554/eLife.57613.

44 Wu, Z., Y. Xiong, S. X. Yu, and D. Lin (2018). “Unsupervised feature learning via non-parametric instance discrimination”. In: Proceedings of the IEEE conference on computer vision and pattern recognition, pp. 3733–3742.

45 Huang, Q., H. He, A. Singh, S.-N. Lim, and A. R. Benson (2020). “Combining label propagation and simple models out-performs graph neural networks”. In: arXiv preprint 2010.13993.

46 Zhu, X. and Z. Ghahramani (2002). “Learning from labeled and unlabeled data with label propagation”. In: ProQuest number: information to all users.

47 Tao, Z., H. Liu, S. Li, Z. Ding, and Y. Fu (2019). “Robust spectral ensemble clustering via rank minimization”. In: ACM Transactions on Knowledge Discovery from Data (TKDD) 13.1, pp. 1–25.

48 Chiu, C.-L., N. Clack, et al. (2022). “Napari: a Python multi-dimensional image viewer platform for the research community”. In: Microscopy and Microanalysis 28.S1, pp. 1576–1577.

49 Yemini, E., A. Lin, A. Nejatbakhsh, E. Varol, R. Sun, G. E. Mena, A. D. Samuel, L. Paninski, V. Venkatachalam, and O. Hobert (2021). “NeuroPAL: a multicolor atlas for whole-brain neuronal identification in C. elegans”. In: Cell 184.1, pp. 272–288.

50 Sharma, A. K., F. Randi, S. Kumar, S. Dvali, and A. M. Leifer (2024). “TWISP: a transgenic worm for interrogating signal propagation in Caenorhabditis elegans”. In: Genetics 227.3, iyae077.

51 Uzel, K., S. Kato, and M. Zimmer (2022). “A set of hub neurons and non-local connectivity features support global brain dynamics in C. elegans”. In: Current Biology 32.16, pp. 3443–3459.

52 Kato, S., H. S. Kaplan, T. Schrödel, S. Skora, T. H. Lindsay, E. Yemini, S. Lockery, and M. Zimmer (Oct. 2015). “Global Brain Dynamics Embed the Motor Command Sequence of Caenorhabditis elegans”. en. In: Cell 163.3, pp. 656–669. issn: 0092-8674. doi: 10.1016/j.cell.2015.09.034.

53 Charles Fieseler (n.d.). Repository for BarlowTrack paper. url: https://github.com/Zimmer-lab/barlow track.

54 Lyons, A. M. and S. Kato (2025). “The tardigrade as an emerging model organism for systems neuroscience”. In: arXiv preprint 2501.06606.

55 Achard, C., T. Kousi, M. Frey, M. Vidal, Y. Paychére, C. Hofmann, A. Iqbal, S. B. Hausmann, S. Pagés, and M. W. Mathis (2025). “CellSeg3D, Self-supervised 3D cell segmentation for fluorescence microscopy”. In: eLife 13, RP99848.

56 He, K., G. Gkioxari, P. Dollár, and R. Girshick (2017). “Mask r-cnn”. In: Proceedings of the IEEE international conference on computer vision, pp. 2961– 2969.

57 Sarlin, P.-E., D. DeTone, T. Malisiewicz, and A. Rabinovich (2020). “SuperGlue: Learning Feature Matching With Graph Neural Networks”. In: pp. 4938–4947.

58 McInnes, L., J. Healy, and J. Melville (2018). “Umap: Uniform manifold approximation and projection for dimension reduction”. In: arXiv preprint 1802.03426.

